# Transcriptomic insights into the role of miR394 in the regulation of flowering time in *Arabidopsis thaliana*

**DOI:** 10.1101/2025.02.15.638417

**Authors:** Federico Belen, Yanel Bernardi, Andrea Reutemann, Abelardo Vegetti, Marcela Dotto

**Affiliations:** Instituto de Ciencias Agropecuarias del Litoral (ICIAGRO L., CONICET-UNL), Facultad de Ciencias Agrarias (UNL), Esperanza, Argentina

**Keywords:** miR394, Arabidopsis, flowering, Transcriptomics

## Abstract

The initiation of flowering is a highly coordinated process that requires synchronization of endogenous plant developmental program with environmental signals through a complex interplay of genetic regulatory networks. The developmental shift from vegetative to reproductive stage is controlled by multiple flowering pathways, which together with other signals participate in the orchestration of flowering time regulation. Collectively, the different pathways converge to modulate the expression of key floral integrator genes, which subsequently trigger the expression of genes related to the differentiation of the vegetative shoot apical meristem into a flower meristem with the subsequent development of floral organs. We had previously demonstrated a role for miR394 in the regulation of flowering time, since *mir394a mir394b* double mutant plants harboring T-DNA insertions in the two *MIR394 Arabidopsis* genes exhibited an early flowering phenotype correlated with modified expression of flowering genes. In the present study we provide transcriptomic data and histological staining of reporter lines to give further insight into the role of miR394 in the regulation of flowering time in *Arabidopsis thaliana* and present an initial bioinformatic characterization of a newly identified lncRNA, which is found partially overlapping the *MIR394B* locus, and therefore we named *<u>MIR</u>394B-<u>AS</u>SOCIATED <u>T</u>RANSCRIPT* (*MIRAST*). We identified differential domains of expression during flower development, which together with the transcriptomic analysis allows us to propose a role for miR394 in the fine tuning of *Arabidopsis* flower development through the regulation of transcription factors and chromatin remodeling factors.

**Key message:** Differential domains of activity of *MIR394A* and *MIR394B* gene promoters together with transcriptomic analysis of mutant plants, indicate that miR394 participates in the fine tuning of *Arabidopsis* flower development through regulation of transcription factors and chromatin remodeling factors.

## Introduction

A plant encounters numerous environmental challenges throughout its lifecycle that must be navigated to ensure survival and reproductive success. As sessile organisms, plants continuously monitor their surroundings, responding to environmental fluctuations through internal regulatory mechanisms that induce specific physiological adaptations (González-Suárez et al. 2023). Among the most critical and intricate of these regulatory mechanisms is the control of flowering time. The initiation of flowering is a highly coordinated process that requires synchronization of the plant developmental program with environmental signals, through a complex interplay of genetic and environmental factors (Bratzel and Turck 2015). In the model plant *Arabidopsis thaliana*, the transition from vegetative growth to reproductive development represents a pivotal stage in the life cycle of the plant, crucial for reproductive success. This developmental shift is controlled by multiple flowering genes included in the photoperiod, vernalization, age, gibberellin and autonomous pathways. Additionally, other signals such as circadian rhythms, sugars and hormones participate in the orchestration of flowering time regulation. Collectively, the different pathways converge to modulate the expression of key floral integrator genes such as *FLOWERING LOCUS T* (*FT*) and *SUPPRESSOR OF OVEREXPRESSION OF CONSTANS 1* (*SOC1*), which subsequently trigger the expression of genes related to the differentiation of the vegetative shoot apical meristem (SAM) into an inflorescence meristem with the subsequent development of floral organs (Amasino 2010; Huijser and Schmid 2011; Liu et al. 2023).

Non-coding RNAs (ncRNAs) have emerged as central regulatory molecules and constitute a significant portion of the transcriptome in eukaryotic organisms. In plants, ncRNAs encompass a diverse array of transcripts, ranging from small ncRNAs, broadly categorized as those with less than 200 nt, to long non-coding RNAs (lncRNAs) longer than 200 nt (Bonnet et al. 2006). Small RNAs include different classes categorized into small interfering RNAs (siRNAs), microRNAs (miRNAs), phased small interfering RNAs (pha-siRNAs) and trans-acting siRNAs (ta-siRNAs), which are typically 20-24-nt long in plants. On the other hand, the classification of lncRNAs is multifaceted, depending on genomic location (intergenic, intronic, or exonic), transcript orientation (sense or antisense), and on their associated functional roles. According to these characteristics, lncRNAs are categorized into long intergenic non-coding RNAs (lincRNAs), intronic non-coding RNAs (incRNAs), promoter-derived antisense RNAs (PAS RNAs) and natural antisense transcripts RNAs (NATs) (Budak et al. 2020; Zhang et al. 2023).

In plants, only a limited number of lncRNAs have been extensively characterized in terms of their functional roles, associated phenotypes, and underlying molecular mechanisms (Jampala et al. 2021). A well-studied subset of these lncRNAs includes those involved in regulating *FLOWERING LOCUS C (FLC)* through epigenetic mechanisms. One such group, collectively referred to as *COOLAIR*, consists of NAT-lncRNAs that fully encompass and repress the transcription of the *FLC* gene via epigenetic modifications (Marquardt et al. 2014; Hawkes et al. 2016). Additionally, *COLDAIR* and the PAS RNA *COLDWRAP* are required for vernalization-mediated epigenetic repression of *FLC*. This repression is driven by the enrichment of Polycomb Repressive Complex 2 (PRC2) and subsequent trimethylation of Histone H3 at Lysine 27 (H3K27me3) at the *FLC* chromatin (Heo and Sung 2011; Kim and Sung 2017; Kim et al. 2017).

On the other hand, microRNAs (miRNAs) represent one of the most extensively studied families of ncRNAs in eukaryotic organisms, playing diverse and critical regulatory roles in both plant and animal systems (Jones-Rhoades et al. 2006; Voinnet 2009; Khraiwesh et al. 2012). miRNAs are transcribed from specific loci, giving rise to a precursor transcript known as primary miRNA (pri-miRNA), subsequently processed into mature miRNAs through several steps involving the RNase III-like enzyme DICER-LIKE 1 (DCL1), among other factors. The mature miRNA is loaded into ARGONAUTE (AGO) proteins to form the RNA-Induced Silencing Complex (RISC) which scans the cytoplasm for complementary target mRNAs, enabling post-transcriptional gene silencing (Bartel 2004; Kurihara and Watanabe 2004; Jones-Rhoades et al. 2006; Voinnet 2009).

Several microRNAs have been identified as crucial regulators of plant growth and development through the post-transcriptional silencing of transcripts coding for transcription factors, which are especially well characterized in *Arabidopsis thaliana*. For instance, miR159 and miR319 have been shown to regulate floral organ development (Rubio-Somoza and Weigel 2013), miR390 participates in auxin signaling (Fahlgren et al. 2006), miR167 regulates auxin response factors (Wu et al. 2006), miR169 plays a role in nitrogen stress responses (Xu et al. 2014) and both miR156 and miR172 are core components of the age-dependent developmental pathway promoting the transition to the adult phase and subsequent flowering (Aukerman and Sakai 2003; Hyun et al. 2017).

A particularly interesting microRNA is miR394, which does not target a transcription factor, but instead regulates the accumulation of the transcript encoding LEAF CURLING RESPONSIVENESS (LCR) protein, part of the F-box family, which are members of SKP1-CULLIN1-FBOX (SCF) ubiquitin-ligases complexes, responsible for marking specific proteins for degradation by the 26S Proteasome (Song et al. 2012). This miRNA has been identified as a significant player in the regulation of various biological processes, including leaf morphology, response to biotic and abiotic stress and development of the shoot apical meristem during embryonic development (Song et al. 2012; Knauer et al. 2013; Song et al. 2013; Song et al. 2016; Tian et al. 2018). In a previous study, we demonstrated a role for miR394 in the regulation of flowering time, since *mir394a mir394b* double mutant plants harboring T-DNA insertions in the two *MIR394 Arabidopsis* genes exhibited an early flowering phenotype correlated with modified expression of flowering genes. Interestingly, our work indicated that the role for miR394 in the regulation of flowering time is independent of LCR and we determined that no additional targets regulated by transcript cleavage for this miRNA. This suggests that additional miR394 targets could exist that would regulate biological processes through alternative molecular mechanisms (Bernardi et al. 2022).

Building on these findings, the present study seeks to further elucidate the role of miR394 in the regulation of flowering time in *Arabidopsis thaliana* and present an initial bioinformatic characterization of a newly identified lncRNA. This lncRNA is found partially overlapping the *MIR394B* locus, and therefore we named it *MIR394B-ASSOCIATED TRANSCRIPT* (*MIRAST*). We used a transcriptomic approach to identify genes with altered expression in *mir394a mir394b* mutant plants, to gain insight into the role of miR394 in the regulation of flowering time and used histological GUS staining to characterize *MIR394A* and *MIR394B* reporter lines. We identified differential domains of expression during flower development, which together with the transcriptomic analysis allows us to propose a role for miR394 in the fine tuning of *Arabidopsis* flower development through the regulation of transcription factors and chromatin remodeling factors.

## Materials and methods

### Plant materials and growth conditions

*Arabidopsis thaliana* accession Columbia-0 (Col-0) was used in this study. The double mutant line in *MIR394* genes, called *mir394a mir394b*, was described by Bernardi et al. (Bernardi et al. 2022). Reporter lines *pMIR394A::GUS* and *pMIR394B::GUS* were generated as described below. For all experiments, seeds were planted in soil, stratified for 3 d at 4 °C and transferred to a growth chamber under long-day conditions (16 h light/8 h dark) at 21-24 °C.

### RNA extraction and RNA sequencing

For transcriptome experiments, aerial part of plants was sampled 14 d after germination, and total RNA was extracted from three biological replicates, being each biological replicate a pool of three plants. Total RNA was isolated from 100 mg of pooled plants using TriPure reagent (Roche), followed by RQ1 DNase (Promega) treatment and posterior purification using the PureLinkTM RNA mini kit (Invitrogen). Poly(A) RNA-Seq sequencing was performed by LC Science (Houston, TX, USA) on strand-specific cDNA libraries, sequenced as 150-bp paired-end reads using Illumina NovaSeq. Sequence reads were deposited in the National Center for Biotechnology Information Sequence Read Archive (NCBI-SRA) under the accession number PRJNA1188835.

### RT-qPCR and stem-loop RT-qPCR

RNA was isolated as described above and first-strand cDNA was obtained by reverse-transcription on 1 μg of DNAse-treated total RNA using ImProm II reverse transcriptase enzyme (Promega). For retrotranscription of mRNA and miR394, oligo(dTN) and stem-loop oligos for miRNA394 were used in the same reaction mix (Varkonyi-Gasic et al. 2007). The obtained cDNA was used as a template for quantitative PCR amplification in a AriaMx 1.6 qPCR System (Agilent), using 1 × *PerfectStart* ® Green qPCR SuperMix, (Transgene Biotech). Primers for each of the genes under study were designed or obtained from previous reports (Additional file 1). Amplifications were performed under the following conditions: 30 s of denaturation at 94 °C, 40 cycles at 94 °C for 5 s, 50 °C for 15 s, and 72°C for 10 s. Three biological replicates and three technical replicates were performed for each sample. Melting curves were determined by measuring fluorescence with increasing temperature (from 65 to 95 °C). Gene expression levels were normalized to that of *Arabidopsis UBIQUITIN10 (UBQ10)*. The efficiency of the primers used for qPCR analysis was determined to be close to 100% for all primers used, and therefore, analysis was performed using the 2^−ΔΔCt^ method (Livak and Schmittgen 2001).

### RNA-seq bioinformatic analysis

A standard bioinformatics pipeline was used to obtain the differentially expressed genes (DEGs) between the two genotypes: FastQC (v 0.12.0, https://www.bioinformatics.babraham.ac.uk/projects/fastqc/) was used to check sequencing data quality, bowtie (v 1.3.1, with default parameters and setting -v 3 option) (Langmead et al. 2009) was used to remove ribosomal RNA, Hisat2 (v 2.2.1, with --rna-strandness option) (Kim et al. 2019) to align reads to the TAIR 10 reference genome (https://arabidopsis.org/download_files/Genes/TAIR10_genome_release/TAIR10_chromosome_files/TAIR10_chr_all.fas.gz) and StringTie (v 2.2.1) (Pertea et al. 2015) for transcript quantification using Araport11 gene models (https://www.arabidopsis.org/download/file?path=Genes/Araport11_genome_release/Araport11_GFF3_genes_transposons.current.gff.gz). Gene-level read count data was generated using the prepDE.py3 script on the General File Format (GFF) files generated by StringTie, and DESeq2 (v 1.36.0) (Love et al. 2014) was used to compute DEGs between Col-0 and *mir394a mir394b.* A threshold of padj ≤ 0.05 and a Fold Change (FC) of ±1.5 was used to identify DEGs. DEGs were further subjected to Gene Ontology (GO) enrichment analysis using AgriGO v2 (Tian et al. 2017), selecting “TAIR genome locus (TAIR10)” as background for the analysis of Biological Processes (BP), Cellular Components (CC) and Molecular Functions (MF).

### Bioinformatic analysis of small RNA-seq data

Small RNA-seq (sRNA-seq) libraries from various tissues and developmental stages of wild-type *Arabidopsis thaliana* ecotype Col-0 were downloaded from NCBI-SRA: 14-day-old seedlings (ERR1937938), seedling roots (SRR20810975), mature leaves (SRR24798075) and flower buds (SRR7760268). FastQC (v 0.12.0) was used for quality control, fastx_clipper from the FASTX Toolkit package (v 0.0.13, http://hannonlab.cshl.edu/fastx_toolkit) was used for adapter trimming, followed by bowtie (v 1.3.1), to remove ribosomal RNA, with -v 3 option and --un to obtain fastq files with clean reads, used next to align to the TAIR10 reference genome using bowtie as well. The aligned reads were visualized using the Integrative Genomics Viewer (v 2.16.2) (Thorvaldsdóttir et al. 2013).

### *MIR394A* and *MIR394B* reporter lines construction

Two reporter lines, named *pMIR394A::GUS* and *pMIR394B::GUS,* were constructed in pKGWFS7 plasmid (Karimi et al. 2002). Fragments of 4423 bp for *MIR394A* and 4628 bp for *MIR394B* were used, corresponding to the upstream regions from the transcription start site of *MIR394A* and *MIR394B* genes. Specific primers for each were used in PCR reactions (Additional File 1) and the sequences of the cloned PCR fragments are reported in Additional File 2. *Arabidopsis thaliana* Col-0 plants were used to generate transgenic reporter lines using the floral dip protocol (Clough and Bent 1998).

### β-glucuronidase (GUS) staining and histochemical analysis

GUS staining protocol was adapted from Jefferson et al, 1987 (Jefferson et al. 1987). Seedlings in the juvenile and adult vegetative stages or inflorescences from *pMIR394A::GUS* and *pMIR394B::GUS* plants were incubated in cold 90% v/v acetone and prefixed for 20 min at 4 °C. The samples were washed for 5 min with a solution containing 100 mM sodium phosphate (pH 7.2), 20 mM EDTA (pH 8), 10% v/v Triton X-100; 0.5 mM potassium ferricyanide and 0.5 mM potassium ferricyanide, followed by an incubation in the same solution containing additional 1.2 mM X-gluc (5-bromo-4-chloro-3-indolyl-β-glucuronide), with vacuum infiltration at 600 mmHg for 45 min. Samples were incubated at 37 °C for 3 days.

For histological sections, the samples were dehydrated in a series of increasing ethanol concentrations (20%, 35%, and 50% v/v) for 30 min each and fixed at room temperature for 30 min in FAA (formaldehyde:acetic acid:ethanol, 5:10:50 % v/v) with gentle agitation at room temperature. Next, a series of incubations was performed under the same conditions, with increasing concentrations of ethanol 70% v/v, 80% v/v and a final overnight incubation in 96% v/v with agitation at 4°C. The samples were clarified by incubating for 30 minutes, under agitation at room temperature with increasing concentrations of Bioclear (Biopack):ethanol mixtures (1:3, 1:1, 3:1). Finally, the samples were incubated overnight in 100% v/v Bioclear with gentle agitation at room temperature. For paraffin embedding the samples were transferred to 100% v/v Bioclear which was gradually replaced with solid modified synthetic paraffin (M.P. 54-57 °C) until the total volume was replaced with new melted paraffin. Plugs were assembled and 13 and 15 µm thick serial cuts were made using a rotary microtome and the sections obtained were mounted with a synthetic substitute of Canada balsam (Canadax®, BIOPUR). All preparations were observed under a Leica DM1000 microscope and photographed using a Canon EOS REBEL T2i digital camera.

## Results

### Characterization of *MIRAST,* a novel long non-coding RNA

To characterize the genomic region containing *MIRAST* and *MIR394B* genes, we analyzed annotations on *Arabidopsis* TAIR10 genome version and identified only the hairpin region of *MIR394B* is annotated between coordinates 28,568,808 - 28,568,928 bp on chromosome 1, but we were able to identify information about the transcription start site of *MIR394B* primary transcript (*pri-MIR394B*) at position 28,568,633, obtained by integrating high-throughput omics data by You et al (You et al. 2017), (Fig. 1A; Table 1). The uncharacterized *MIRAST* was annotated on the complementary strand by the Arapor11 project and was classified as an intergenic lncRNA (Cheng et al. 2017a). Hence, combining the gathered information, here we show that these transcripts present a 45 bp overlap, indicating *MIRAST* could also be classified as a natural antisense transcript (NAT)-lncRNA or promoter-derived antisense RNAs (PAS RNAs) (Fig. 1, Table 1).

**Fig. 1:**
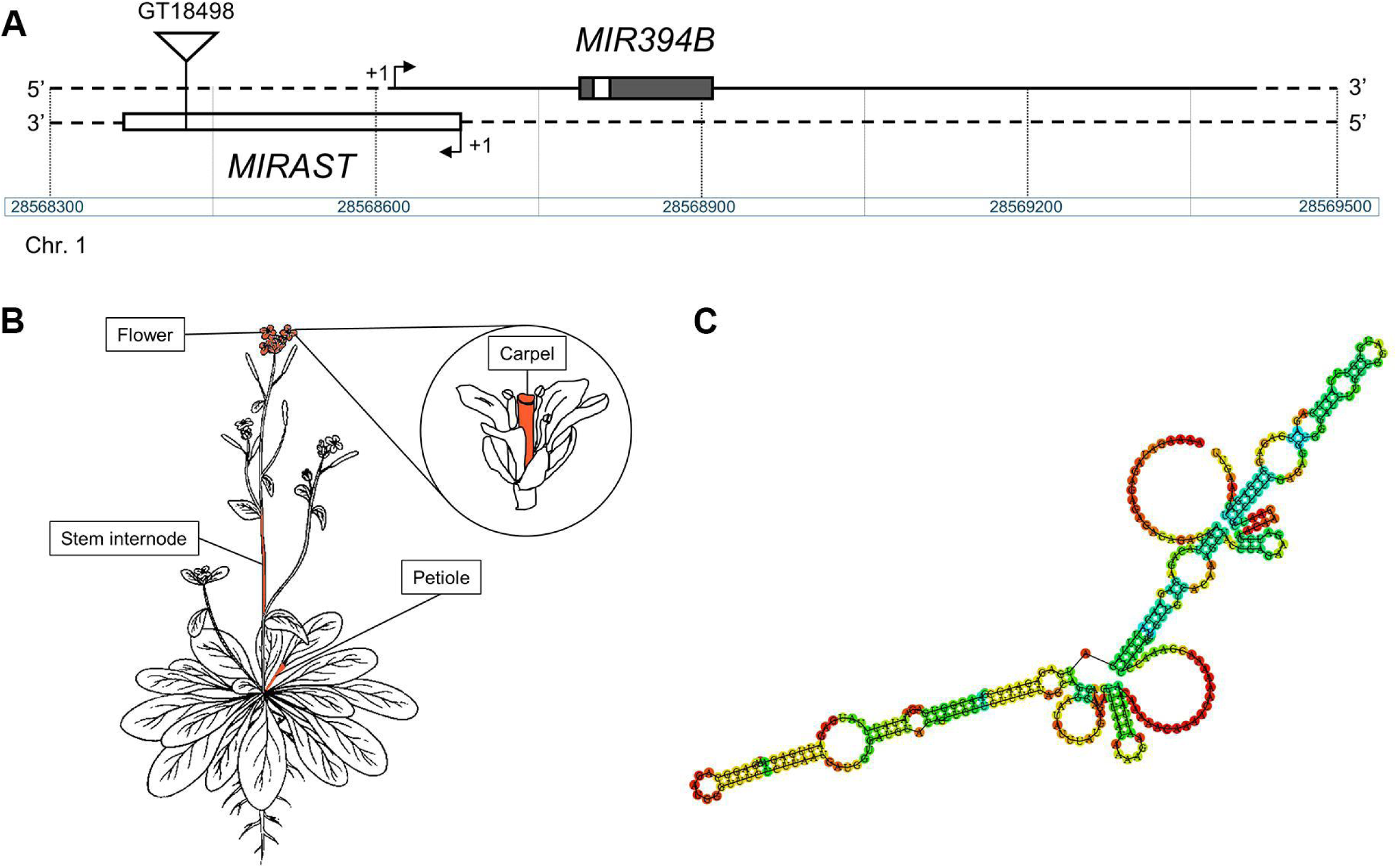
(A) Genomic region containing *MIR394B* and *MIRAST* genes. Insertion in GT18498 line is indicated with a triangle (not to scale). Arrows indicate transcription start site (+1). White rectangle in *MIR394B* representation corresponds to the mature miR394 location. (B) Specificity of expression according to τ (Tau) index > 0.54 (see text for details). (C) Predicted secondary structure of *MIRAST* using RNA-fold.

**Table 1.** Coordinates of *MIR394B* and *MIRAST*.

| Gene | Coordinates |
| --- | --- |
| *pri-MIR394B* | chr1: 28568633 – 28569416 |
| Hairpin *MIR394B* | chr1: 28568808 – 28568928 |
| *MIRAST* | chr1: 28568367 - 28568677 |

Previously, we characterized double mutant plants harboring a knock-out insertional mutation in *MIR394A* gene (GK491D11) and a knock-down insertional mutation in *MIR394B* gene promoter (GT18498), which results in a 5-fold decrease in mature miR394 accumulation (Bernardi et al. 2022). Taking into consideration the genomic annotations presented in Figure 1, we established that insertion in GT18498 mutant line is located within *MIRAST*, which overlaps with the promoter region of *MIR394B* gene, (Fig. 1A) (Bernardi et al. 2022).

Next, we used publicly available expression data to gain insight into *MIRAST* expression pattern (Klepikova et al. 2016; Corona-Gomez et al. 2022). This lncRNA exhibits a low global expression, with specific accumulation in seeds, leaf petioles, carpels and stem internode according to Klepikova’s Atlas (Suppl. Fig. S1). To further assess the tissue specificity of *MIRAST*, we used previously developed Tau index, which indicates gene expression specificity. Tau values range from 0, representing housekeeping genes, to 1, representing genes that are exclusively expressed in a single tissue (Yanai et al. 2005). Information for *MIRAST* was extracted from Corona-Gomez *et. al.* (2022), where genes with τ values higher than the median τ of mRNAs (0.54) are considered tissue-specific or developmental stage-specific. The τ index calculation for *MIRAST* confirmed its specific expression in petiole (τ = 0.627), stem internode (τ = 0.783), whole flower (τ = 0.586) and carpel (τ = 0.759) (Fig. 1B), reinforcing its potential functional relevance in these particular tissues and developmental contexts involving these organs and plant tissues.

Secondary structure prediction indicates *MIRAST* folds into a complex structure with two long arms containing hairpin-folded regions that could be processed by DICER-LIKE enzymes (Fig. 1C). Therefore, we used small RNA sequencing data from public databases to analyze whether this lncRNA could be a source of siRNAs. We included information from 14-day-old seedlings, seedling roots, mature leaves and flower buds (Additional File 3). In addition, we took into consideration that *MIRAST* and *pri-MIR394B* transcripts have complementary sequences in their 5’ regions (Fig. 1A), being possible for them to hybridize and generate a dsRNA region when they are co-expressed. Observation of mapped reads in the genomic region containing these two genes shows that mature miR394 is the most abundant small RNA derived from this genomic region (Suppl. Fig. S2). In addition, even though a few reads map to the overlapping region, a detailed analysis indicates they are between 26 and 32 nucleotides-long (Suppl. Fig. S2), which is longer than typical plant small RNAs, and no siRNAs were detected that would derive from the hairpin structures predicted for *MIRAST*. Hence, we conclude that this lncRNA is not likely to function as a source of small RNAs produced by hairpin-like structures or through the formation of dsRNA processed by DCL enzymes.

### Transcriptome analysis in *mir394a mir394b* mutant plants

In a previous initial characterization of *mir394a mir394b* double mutant plants, we showed they express 5-fold less mature miR394 than Col-0 plants (Fig. 2A) and present an early flowering phenotype, which is independent of LCR (Bernardi et al. 2022). To gain insight into the regulatory functions of miR394 and to analyze a possible function for the uncharacterized *MIRAST*, we used RNA-seq to globally analyze gene expression in *mir394a mir394b* mutant plants. Even though we now know the mutation in *MIR394B* also affects the gene coding for *MIRAST* (Fig. 1A), we maintain the name of this mutant line to be consistent with our previous publication (Bernardi et al. 2022). Transcriptome analysis was conducted on 14-days-old plants, grown under long-day conditions, since plants at this stage are typically transitioning to reproductive development (Boyes et al. 2001; Klepikova et al. 2016) (Fig. 2A). To identify the differentially expressed genes (DEGs) between the *mir394a mir394b* mutant plants and wild-type Col-0 plants, a standard bioinformatic pipeline was used to process and analyze the sequencing data, yielding approximately 17 million uniquely mapped reads from each sample and Pearson r^2^ correlation values between 0.95 and 0.99, indicating consistency of the data and allowing for comparisons using all three biological replicates (Additional file 3). Analysis of expression values for *MIRAST* allowed us to confirm that this gene has a non-detectable expression in Col-0 plants, with no differential expression in *mir394a mir394b* mutant plants (Fig. 2A). Therefore, we continued our transcriptomic analysis considering that mutations in these plants mainly affect miR394 action and a functional characterization of *MIRAST* would not be possible using this transcriptomic approach.

**Fig. 2:**
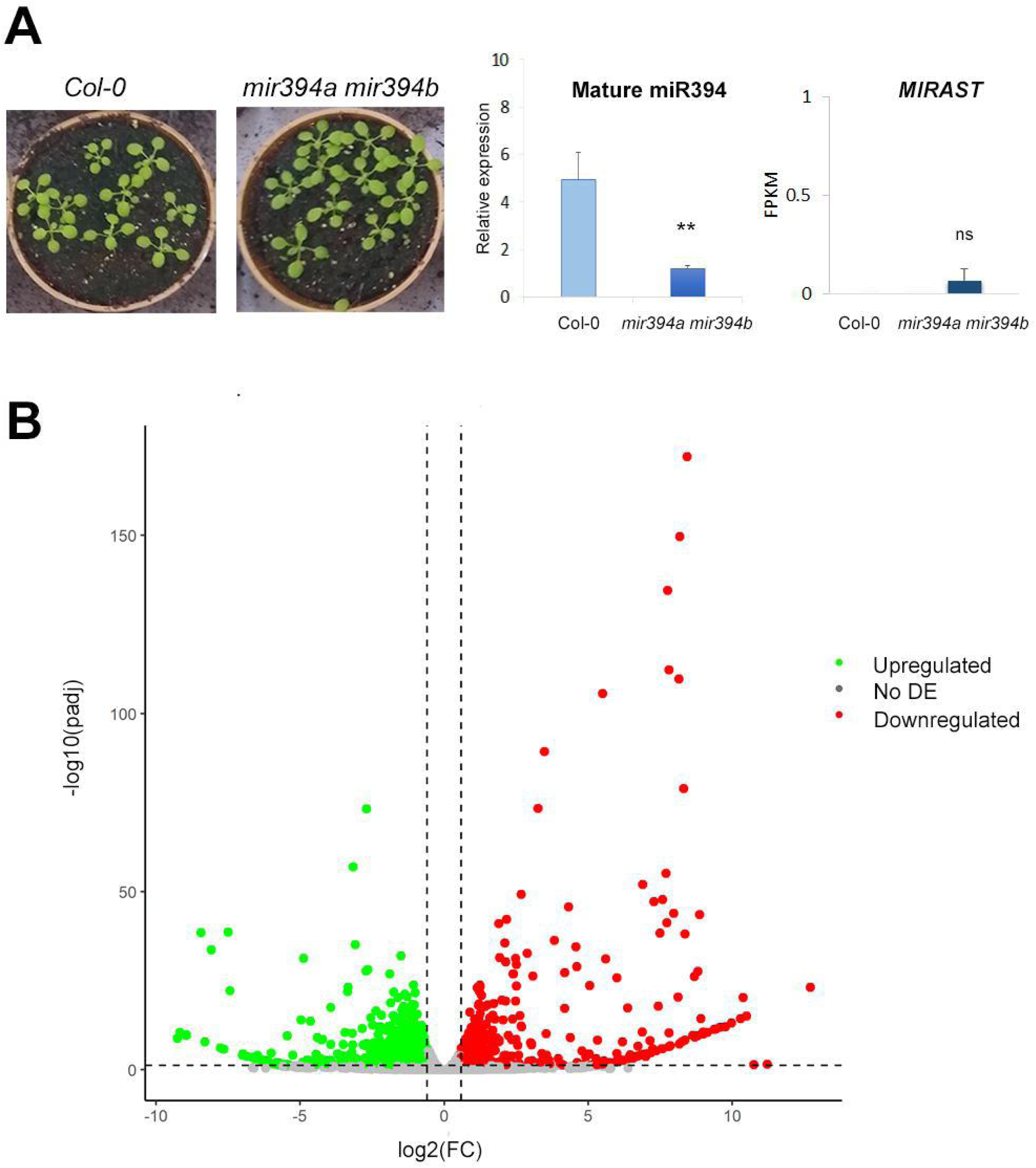
A) Plants grown under long-day conditions for 14 days, used for transcriptome analysis, and expression of *MIRAST*. B) Volcano plot displaying DEGs: 1349 upregulated genes (green dots) and 1056 downregulated genes (red dots) in *mir394a mir394b* compared to Col-0. Gray dots correspond to not differentially expressed genes (No DE) according to padj < 0.005 and FC (Col-0/*mir394a mir394b* or *mir394a mir394b/Col-0)* ≥ 1.5.

Our results identified a total of 24691 genes expressed in the analyzed samples, with a total of 2405 DEGs, of which 1349 were upregulated and 1056 were downregulated in *mir394a mir394b* mutants compared to Col-0 (Fig. 2B; Additional file 4).

### Gene Ontology enrichment analysis of differentially expressed genes

To gain insight into the processes affected in the mutant plants, we used GO Term Enrichment analysis for classification of DEGs. A total of 423 enriched terms at different hierarchical levels were identified, among which 348 correspond to the category Biological Process (BP), 18 to Molecular Function (MF) and 57 to Cellular Component (CC), (Additional File 5). A summary of GO terms in the three categories is presented in Fig. 3A, showing the first enriched term within each top level. The enriched terms within the Biological Process category encompasses a broad range of processes, including terms related to the interaction with the environment, such as ‘response to stimulus’ and ‘immune system process’, as well as a number of terms related to signaling and regulation of different aspects of plant growth and development, such as ‘regulation of biological process’ and ‘biological regulation’, ‘ribonucleoprotein complex biogenesis’, ‘signaling’, ‘death’, ‘post-embryonic development’ and ‘floral whorl development’. Within Cellular Components, the terms ‘cell’ and ‘organelle’ were found enriched, which include enriched downstream terms related to chloroplasts and chloroplast parts (Additional file 5). In the category Molecular Function, one of the enriched terms is related to gene expression (‘transcriptional regulator activity’), being ‘binding’ and ‘catalytic activity’ the most significantly enriched terms (Fig. 3A). The broad aspects encompassed by the enriched terms seem to be in agreement with the several different functions that have been described for miR394, some of which have been described in detail (Song et al. 2012; Knauer et al. 2013; Song et al. 2013; Song et al. 2016; Zhang et al. 2021; Lu et al. 2023).

**Fig. 3:**
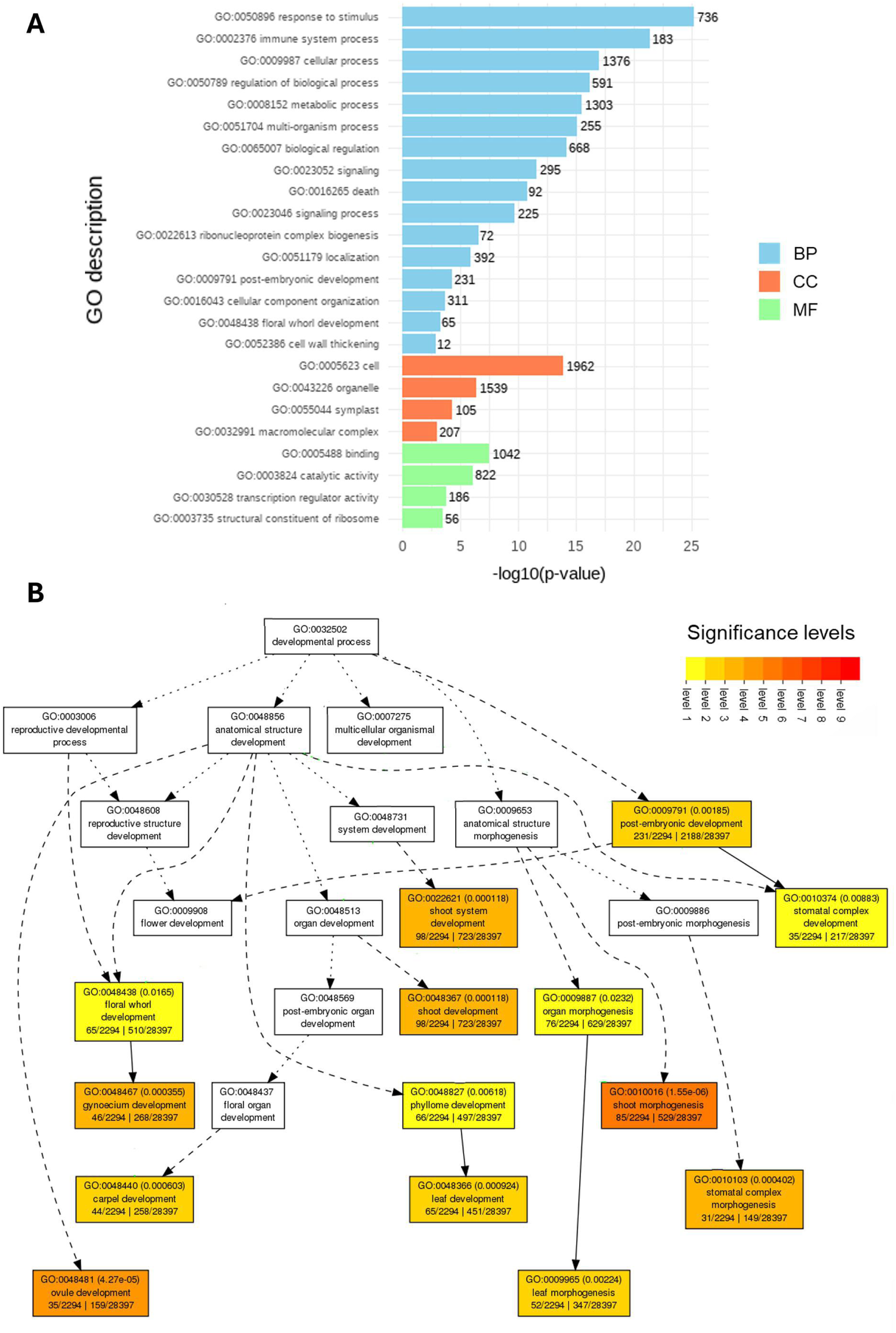
Gene Ontology (GO) enrichment analysis of differentially expressed genes in *mir394a mir394b* double mutant plants. (A) Bar chart showing enriched GO terms categorized in Biological Process (BP, blue), Cellular Component (CC, orange), and Molecular Function (MF, green). Numbers next to bars indicate the number of DEGs for each term. (B) Hierarchical representation of enriched GO terms related to developmental processes. Boxes represent individual GO terms, with the FDR value indicated between brackets for enriched terms and different colors indicating significance levels, from yellow (level 1) to red (level 9). Dashed lines show parent/child relationships between terms.

Taking into consideration our previous publication showing a role for miR394 in the regulation of flowering time (Bernardi et al. 2022), we explored the enriched GO terms related to the term ‘developmental processes’. The enriched child terms shown in Fig. 3B are related to developmental pathways affected in the mutant plants that have been shown to be regulated by miR394. For example, the enriched terms ‘shoot system development’, ‘leaf development’ and ‘leaf morphogenesis’ are related to miR394 functions already explored in previous works (Song et al. 2012; Knauer et al. 2013). In addition, several enriched terms are associated with flowering, such as ‘floral whorl development’ and those related to the development of the female reproductive structures (gynoecium, carpel and ovule), which are of interest to gain insight into the role of miR394 in the regulation of flowering time described in our previous work (Bernardi et al. 2022).

### Differential expression of flowering genes

Because of the early flowering phenotype observed in *mir394a mir394b* plants (Bernardi et al. 2022) and the identification of enriched terms related to flower development in the GO term enrichment analysis (Fig. 3), we focused the following analysis on the molecular pathways regulating flowering time and flower development. First, we analyzed flowering genes typically categorized into the main flowering pathways: age, autonomous, vernalization, thermosensory, photoperiod and gibberellin (Additional File 6) (Wu et al. 2009; Wang 2014; Fernández et al. 2016; Cheng et al. 2017b; Leijten et al. 2018; Zheng et al. 2019; Sharma et al. 2020; Kim 2020; Qi et al. 2022). Among 49 analyzed flowering genes, only 3 *SPL* genes showed differential expression between Col-0 and mutant plants (Additional File 6). Both *SPL2* and *SPL11* transcripts are 2-fold and 1.5-fold downregulated in mutant plants, respectively, whereas *SPL3* expression is 2.2-fold upregulated. The rest of the miR156-regulated *SPL* genes did not show differential expression and neither did the miR172-regulated genes categorized in the age pathway (Additional File 6). Moreover, none of the analyzed genes from the autonomous and vernalization pathways showed differential expression, as well as the thermosensory, photoperiod and gibberellin genes (Additional File 6). These results suggest that the mutations in these plants do not affect most of the regulatory networks converging at floral integrator genes *SOC1* and *FT*.

Next, we analyzed floral integrators and observed that *SOC1* expression is upregulated in the mutant plants, which is in accordance with the early flowering phenotype described for these mutants (Bernardi et al. 2022); nonetheless, *FT* expression remains unchanged between both genotypes (Fig. 4A; Additional File 7). Higher *SOC1* expression indicates that mutant plants have probably already transitioned to reproductive development at the sampling point. Hence, when we analyzed meristem identity genes, we observed a 2.4-fold increase for the flower meristem identity gene *FUL* in double mutant plants, while *LFY* expression, another inflorescence meristem identity gene, remains unchanged in mutant plants (Fig. 4A, Additional File 7). We continue to analyze the expression of genes involved in flower development and homeotic genes from the ABCDE model, identifying several genes with differential expression in the mutant plants (Fig. 4B-C; Additional File 7). *HUA ENHANCER 4 (HEN4),* which regulates stamen development, is downregulated in mutant plants, as well as *NGATHA1 (NGA1)* and *NGATHA3 (NGA3),* involved in style development (Fig. 4B; Additional File 7). Conversely, *REPRODUCTIVE MERISTEM 22 (REM22)* and *TGA MOTIF-BINDING PROTEIN 9 (TGA9)* are upregulated in mutant plants, which participate in carpel and anther development, respectively (Fig. 4B; Additional File 7). Neither *STYLISH 1 (STY1)* nor *STYLISH2 (STY2)*, involved in stylo development, were differentially expressed between Col-0 and mutant plants (Fig. 4B, Additional File 7).

**Fig. 4:**
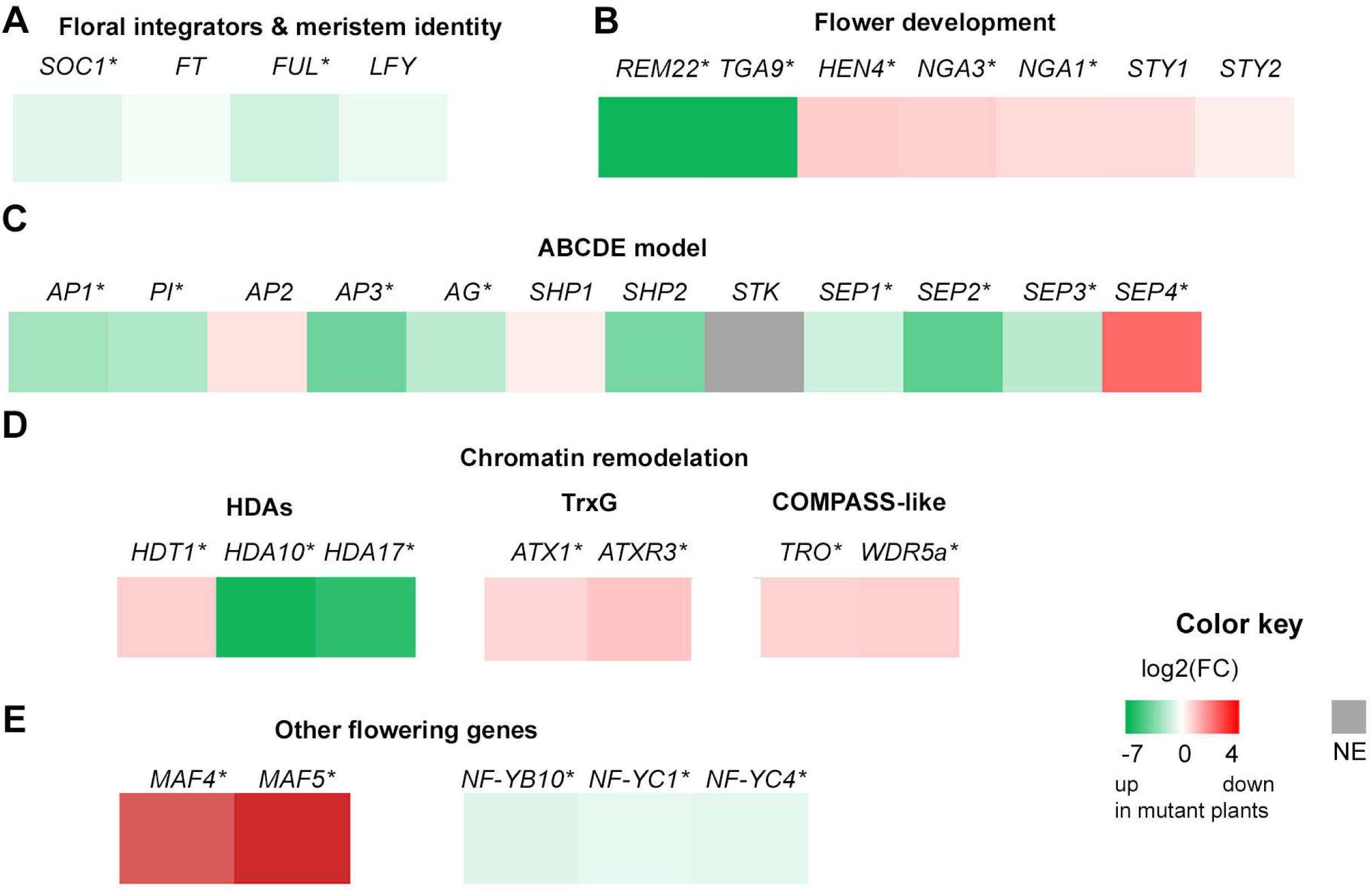
Expression of flowering genes in Col-0 and *mir394a mir394b* plants. A) Expression of floral integrators and inflorescence meristem identity genes. B) Expression of genes involved in flower development. C) Expression of genes of the ABCDE model. D) Expression of genes involved in chromatin remodeling: Histone deacetylases, genes from Thritorax (TrxG) and COMPASS-like complexes. E) Expression of other genes involved in the regulation of flowering time. Log2 is represented for the fold-change (FC) *Col-0/mir394a mir394b*. Green and red indicate genes upregulated and downregulated in mutant plants, respectively. White corresponds to genes unchanged between both samples and gray indicates gene is not expressed (NE). Asterisk next to the gene name indicates differential expression between samples. See text for details.

When we analyzed genes from the ABCDE model, we established that most of them are differentially expressed between the compared genotypes (Fig. 4C). Expression of both *AP1* and *PI* (class A genes) is upregulated in mutant plants, as well as *AP3* (class B) and *AG* (class C), (Fig. 4C, Additional File 7). On the other hand, neither *SHP1* nor *SHP2* (class D genes) are differentially expressed and *STK* is not expressed at this point (Fig. 4C, gray color). Finally, *SEPALLATA* genes (class E) are differentially expressed between samples, being *SEP1*, *SEP2* and *SEP3* upregulated and *SEP4* downregulated in mutant plants (Fig. 4C, Additional File 7). These changes in the expression of flowering genes are in agreement with the early flowering phenotype of mutant plants, indicating floral organ formation is advanced in comparison to Col-0 plants.

Considering regulation of flowering is subject to epigenetic changes in some of the differentially expressed genes mentioned above, such as the floral integrator *SOC1* and homeotic genes like *AP3* and *AG*, among others (Guo et al. 2015) we analyzed expression changes for chromatin remodeling factors. We identified *HISTONE DEACETYLASE 10 (HDA10)* and *HDA17* are upregulated in mutant plants, while *HDT1 (HDA3)* expression is higher in Col-0 plants (Fig. 4D, Additional File 7). Analysis of genes involved in histone methylation from POLYCOMB REPRESSIVE COMPLEX 1 and 2 showed that they are not differentially expressed in the mutant plants (Additional File 7). Interestingly, we identified Thritorax Group factors that are also differentially expressed in mutant plants (Fig. 4D). These genes participate in H3K4 methylation, an activating mark, and have been shown to be involved in the formation of flower organs and play a role in the regulation of homeotic genes (Jiang et al. 2011; Fromm and Avramova 2014; Chen et al. 2017). We identified that *ATX1* and *ATRX3/SDG2,* as well as *TRO*, *WDR5a*, which encode components of a COMPASS-like complex, are downregulated in mutant plants (Fig. 4D, Additional File 7), suggesting a link between miR394 action and chromatin remodeling factors to regulate flowering in *Arabidopsis*.

Further exploration of DEGs allowed us to identify additional genes that have been reported to play roles in the regulation of flowering time (Fig. 4E, Additional File 7), which include *MAF4* and *MAF5*, two members of a family of FLC-related MADS-box transcription factors (Yoo et al. 2011; Gu et al. 2013), as well as three members of the Nuclear Factor Y family of transcription factors, NF-YB10, NF-YC1 and NF-YC4 (Kumimoto et al. 2008; Siriwardana et al. 2016; Zhao et al. 2017).

Taken together, the analysis of DEGs related to the regulation of flowering time and flower development, indicates that *mir394a mir394b* plants had transitioned to reproductive stage at the sampling point, and this occurred earlier than for Col-0 plants. The transcriptomic changes indicate that miR394 affects the fine tuning of flower development through the action of several transcription factors and chromatin remodeling factors.

### GUS staining of *MIR394A* and *MIR394B* reporter lines

Taking into consideration the differentially expressed genes related to flower development and floral organ formation, we analyzed the domains of expression of *MIR394A* and *MIR394B* genes through the generation of reporter lines *pMIR394A::GUS* and *pMIR394B::GUS*. First, staining was performed on 14-days-old whole seedlings in the vegetative stage (Fig. 5) and we also conducted a detailed analysis on young and mature flowers of 30-days-old plants (Fig. 6). In addition, histological sections of the stained tissues were prepared (Fig. 5C, F; Fig. 7).

**Fig. 5:**
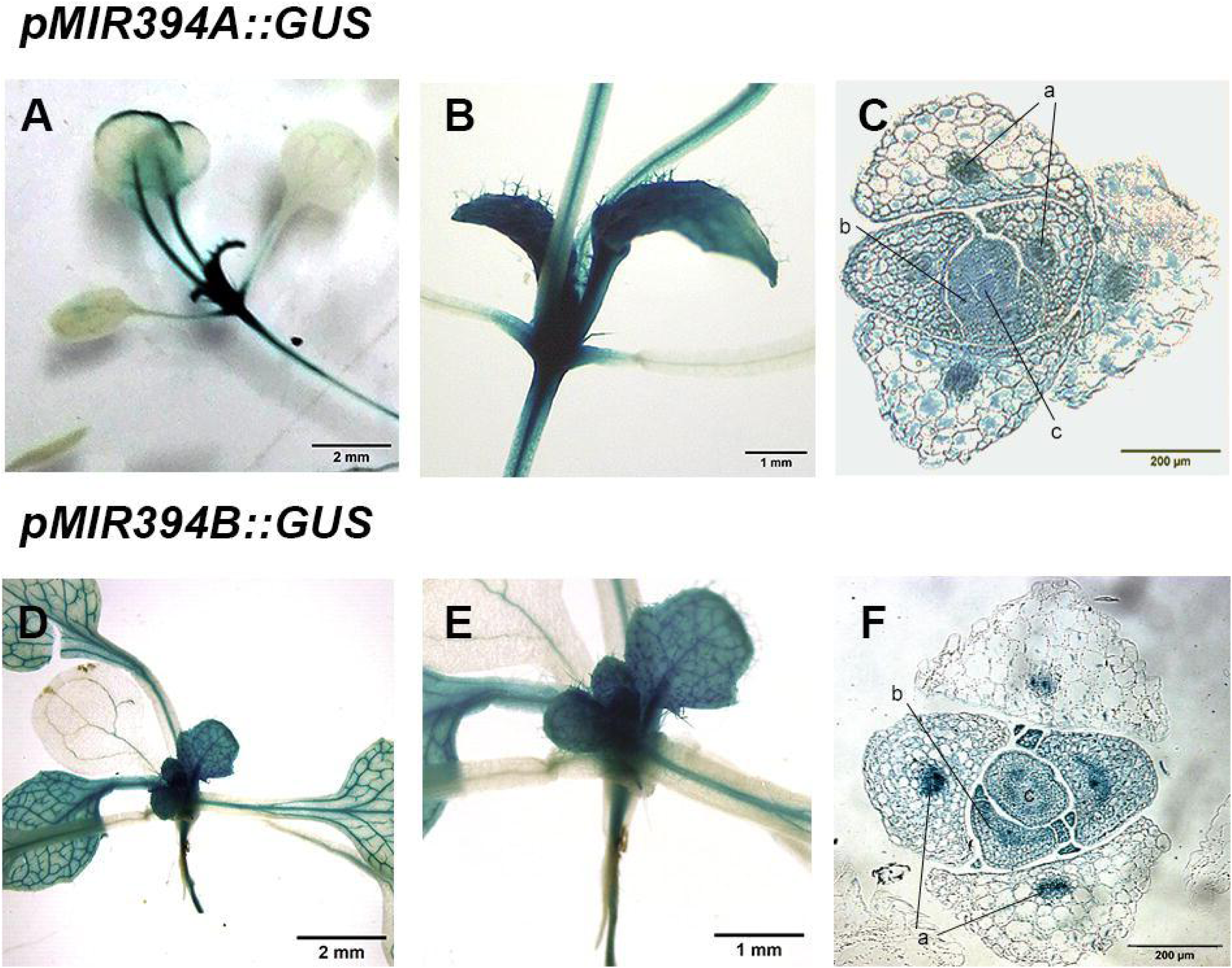
Histochemical GUS staining of *pMIR394A::GUS* and *pMIR394B::GUS* plants. (A, D) 14-days-old plants; (B, E) enlargement of the shoot apical region in A and D; (C, F) 15 μm thick transversal section of SAM. Reference: (a) vascular bundle; (b) leaf primordium; (c) SAM.

**Fig. 6:**
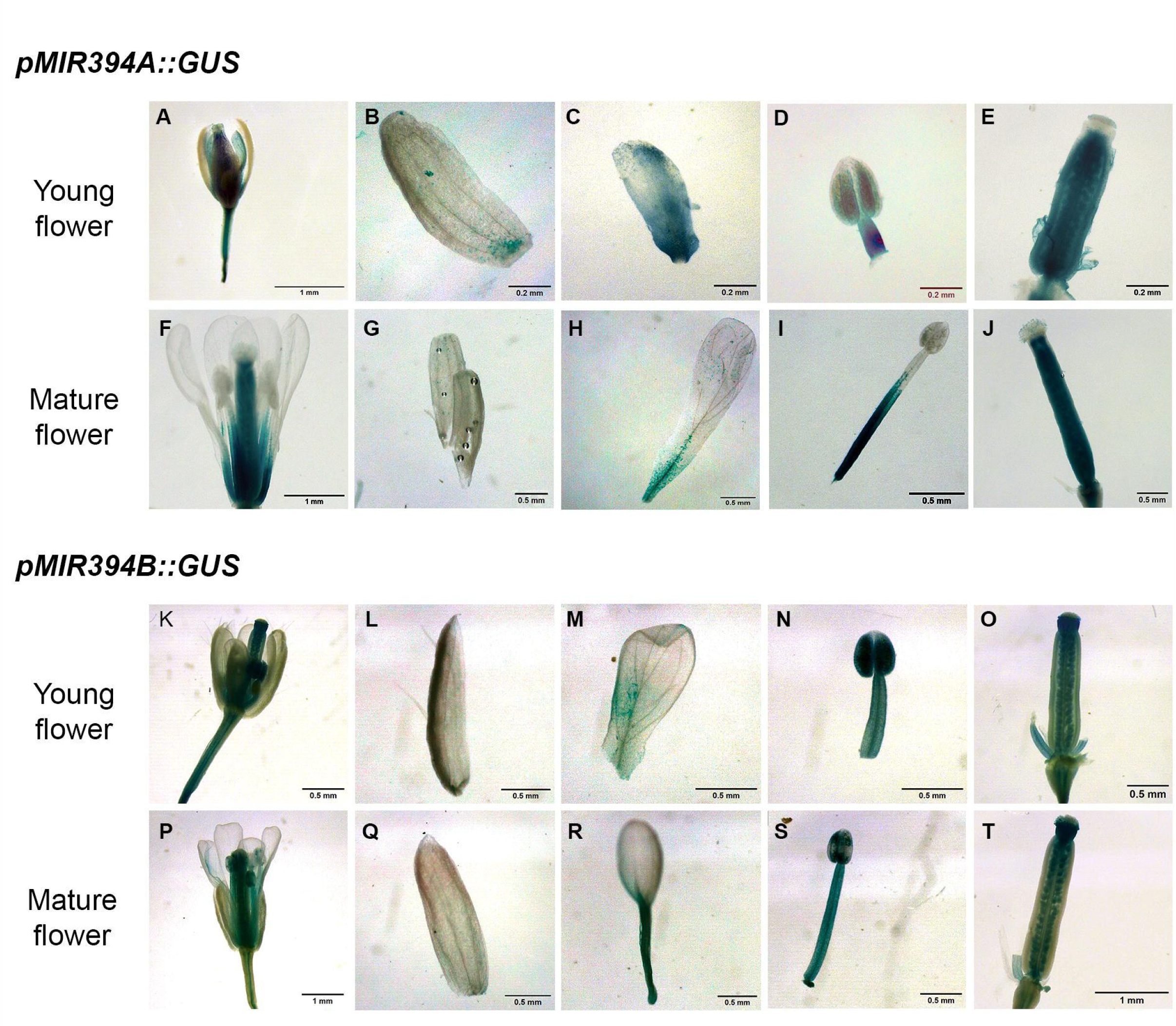
GUS staining of young and mature flowers in 30-days-old plants from *pMIR394A::GUS* reporter line (A-J) and *pMIR394B::GUS* reporter line (K-T). (A, K) Young flowers; (F, P) mature flowers; (B, G, L, Q) sepals; (C, H, M, R) petals; (D, I, N, S) stamen filament and anther; (E, J, O, T) gynoecium.

**Fig. 7:**
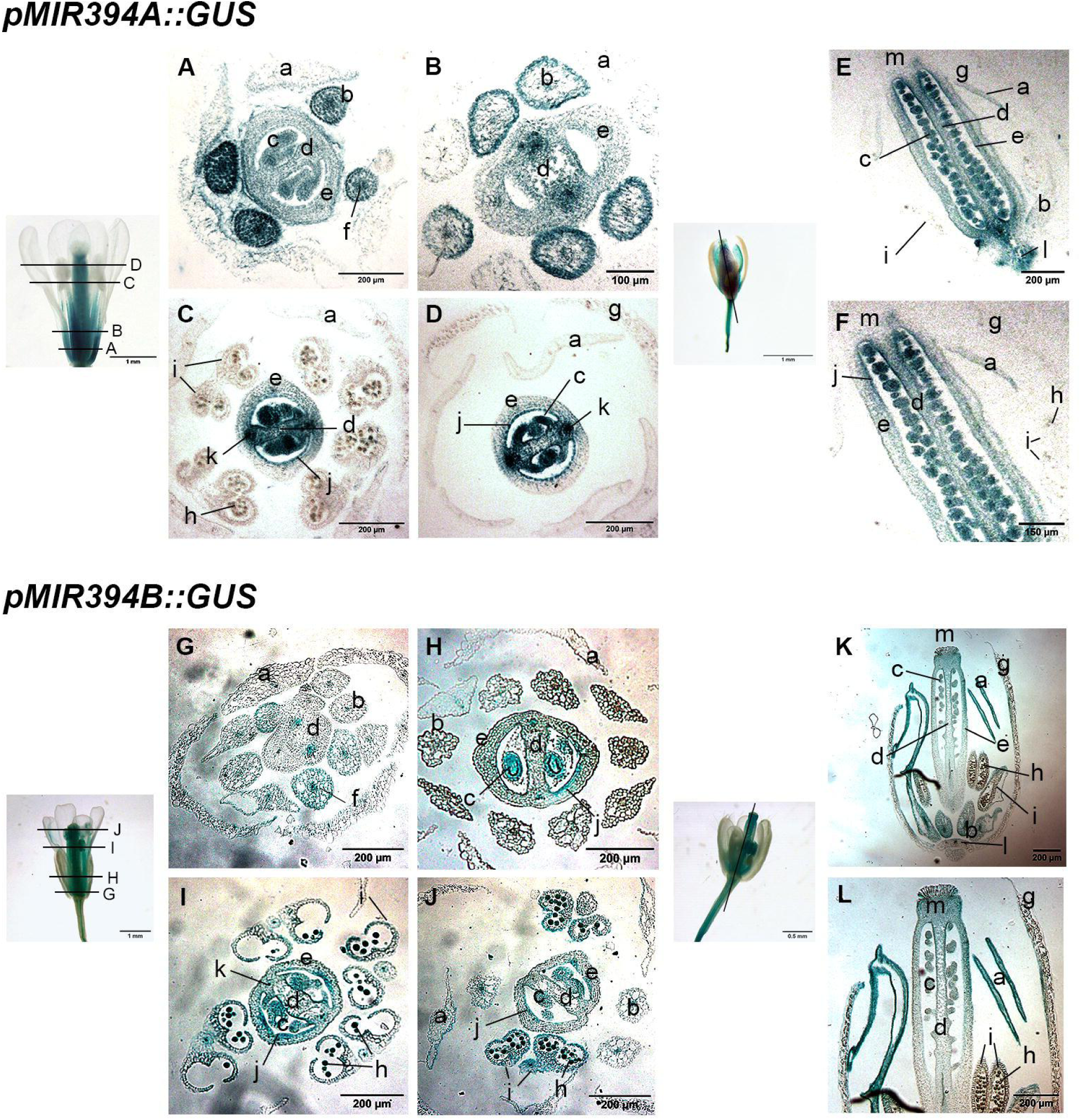
Histochemical GUS staining of flowers for *pMIR394A::GUS* and *pMIR394B::GUS* plants. (A-D and G-J) Cross sections corresponding to the four zones indicated in the mature flower on the left; (E, K) longitudinal sections indicated with a line on the young flowers; (F, L) enlargement of the upper region shown in E and K. (a) Petals. (b) Stamen filaments. (c) Seminal primordium. (d) Carpelar septum. (e) Carpels. (f) Filament vascular tissue. (g) Sepals. (h) Pollen grain. (i) Anther. (j) Adaxial epidermis of carpels. (k) Vascular bundle. (l) Receptacle. (m) Stigma.

GUS staining indicated that the *MIR394A* and *MIR394B* gene promoters were highly active in the shoot apex of seedlings (Fig. 5A, D). Close-up images of these regions revealed intense staining in the shoot apical regions and in young developing leaves (Fig. 5B, E). However, in older leaves *GUS* expression is limited to petioles, main nerves and leaf margins for *MIR394A* promoter (Fig. 5A), whereas the *MIR394B* gene promoter is active in leaf veins (Fig. 5D). Cross sections of the shoot apical region of 14-days-old plants indicate that both reporter lines show strong staining in vascular bundles (a), leaf primordia (b) and the shoot apical meristem (SAM) (c), (Fig. 5C, F).

In young flowers, the expression of *MIR394A* is prevalent in the area closest to the receptacle in sepals (Fig. 6A, B), the basal area in petals (Fig. 6C), in the stamen filaments but not in anthers (Fig. 6D) and the entire gynoecium, excluding stigmas (Fig. E). On the other hand, in young flowers (Fig. 6K), No staining was observed in sepals for *MIR394B* reporter line (Fig. 6L). The *MIR394B* promoter exhibits activity in the mid-petal area (Fig. 6M), complete staining in anthers, but in the stamen ilament activity appears restricted to the vasculature (Fig. 6N). Another difference resides in the gynoecium, in which activity for *MIR394B* promoter is stronger in seed primordia and stigmas (Fig. 6O).

In mature flowers (Fig. F, P) the GUS staining in sepals disappears entirely in both reporter lines (Fig. 6G, Q), whereas in petals, it is localized in vascular tissues close to the receptacle for both reporter lines (Fig. 6H, R). The activity of the *MIR394A* promoter is higher in the basal zone of the stamen filaments and decreases towards the zone near the anthers (Fig. 6I), whereas the activity of the *MIR394B* promoter is high and uniform throughout stamen filaments and anthers (Fig. 6S). Furthermore, the staining for both promoters in the gynoecium maintains the pattern observed in young flowers (Fig. 6J, T).

Furthermore, to study in detail the domains of activity of these two gene promoters, we analyzed both cross histological sections of mature flowers and longitudinal sections for young flowers (Fig. 7). In mature flowers, the *MIR394A* promoter is active in the adaxial part of petals (a), near the receptacle (Fig. 7A), but is not active in the distal part of petals (Fig. 7C-D), whereas the *MIR394B* promoter is not active in petals of mature flowers (Fig. 7G-J), but staining was detected in petals in young flowers (Fig. 7K-L). *MIR394A* promoter shows higher activity in the basal part of stamens (b) and in the filaments of stamens (f), (Fig. 7A-B), whereas *MIR394B* activity was detected only in the filaments of stamens (Fig. 7G-H). In addition, activity of *MIR394B* promoter is observed in anthers (i) and pollen grains (h), (Fig. 7I-J), whereas the *MIR394A* promoter did not show activity in either of them (Fig. 7C). Regarding expression patterns common to both, they show activity in seminal primordia (c), carpelar septum (d) and carpels (e), (Fig. 7A-D, G-J), with *MIR394A* presenting stronger staining intensity in these three. In addition, *GUS* expression is localized in the adaxial epidermis of fused carpels (j) for both promoters (Fig. 7C-D, I-J).

Longitudinal sections of young flowers reveal *MIR394A* promoter activity in the region closest to the receptacle (l), as well as in petals (a), seminal primordia (c), carpelar septum (d), carpels (e) and adaxial epidermis of carpels (j), (Figure 7E-F). *MIR394B* promoter is active in petals (a), stamen filaments (b), seminal primordia (c) and stigmas (m), but no activity is detected in developing pollen grains (h), (Figure 7K-L).

Taken together, these results indicate that *MIR394A* and *MIR394B* genes are expressed in vegetative and reproductive stages, with differential patterns of expression that may contribute to the fine tuning of the development of vegetative organs and flowers.

## Discussion

The results presented here allowed us to introduce an initial characterization of a novel lncRNA, genomically associated to *MIR394B* gene, which we have named *MIRAST: <u>MIR</u>394B <u>AS</u>sociated <u>T</u>ranscript*. However, our transcriptomic analysis in combination with existing expression data indicates that *MIRAST* expression is barely detected in our samples, either Col-0 or *mir394a mir394b* mutants. Therefore, the transcriptomic analysis focused on getting a deeper understanding of the role of miR394 in the regulation of flowering time to complement our previous study presenting an initial characterization of the early flowering phenotype observed in *mir394a mir394b Arabidopsis* mutant plants (Bernardi et al. 2022).

Here we show that mutations in miR394 affect flowering time through deregulation of several transcription factors and chromatin remodeling factors that contribute to the establishment of flowers and floral organ development. In line with our previous work characterizing the early flowering phenotype of *mir394a mir394b* mutants (Bernardi et al. 2022), the transcriptomic data indicates that mutant plants present an earlier transition to reproductive development in comparison to Col-0 plants. Our current analysis indicates that mutant plants had already transitioned to reproductive stage at the sampling point and the flower meristem had been established. Higher expression in mutant plants of the floral integrator *SOC1* and the floral meristem identity gene *FUL* supports this conclusion. Evidence of an advanced flower formation stage in mutant plants resided in the expression of flower organ identity genes, responsible for specification of the four whorls, which are upregulated in comparison to Col-0 plants: *PI* and *AP1* (class A), *AP3* (class B), *AG* (class C) and *SEP1*, *SEP2* and *SEP3* (class E). However, no expression for class D genes (*SHP1*, *SHP2* and *STK*) was detected, which is consistent with these genes being regulated by ABC genes and will probably be activated at later stages (Alvarez-Buylla et al. 2010). Conversely, *SEP4* showed lower expression than Col-0 plants, which could be explained by the fact that this gene has a particular pattern of expression, starting earlier than the rest of the *SEPALLATA* genes, has a broader pattern being also expressed in leaves and stems and considered also a meristem identity gene, marking clear differences with *SEP1*, *SEP2* and *SEP3* (Ma et al. 1991; Ditta et al. 2004; Jetha et al. 2014). Finally, genes involved in the formation of other parts of the plant, including ovary, anther and style (*REM22, TGA9, HEN4, NGA1* and *NGA3*) (Trigueros et al. 2009; Ge et al. 2010; Romanel et al. 2011; Wils and Kaufmann 2017) are also deregulated in mutant plants, revealing that a link between miR394 and the regulatory networks controlling flower organ development.

Interestingly, chromatin remodeling factors are also affected in mutant plants. Epigenetic mechanisms include post-translational histone modifications, especially histone methylation and acetylation/deacetylation that play essential roles in plant development and plasticity (Baulcombe and Dean 2014; Zhao et al. 2019). Histone acetylation is a modification that associates to active genetic loci. There are multiple histone acetyltransferases (HATs) and histone deacetylases (HDAs) that add or remove acetyl groups on histones, respectively, several of which are expressed in inflorescences and floral meristems, and have also been reported to participate in the regulation of flowering genes (Pandey et al. 2002; Hollender and Liu 2008). This study shows that three histone deacetylases present altered expression in mutant plants, specifically *HDT1/HDA3, HDA10* and *HDA17*. Modifications involving histone methylation include the action of protein complexes like the Trithorax (TrxG) and Polycomb (PcG) groups which are the two most important complexes that regulate gene expression at the chromatin level through the deposition of H3K4me3 and H3K36me3 activation marks (TrxG), and H3K27me3 repressive marks (PcG), (Bemer and Grossniklaus 2012; de la Paz Sanchez et al. 2015; Pu and Sung 2015). We did not observe differences in genes coding for the PcG complexes PRC1 and PRC2 in our study, which have been extensively studied and documented in plants (Kuwabara and Gruissem 2014; Avramova 2015; Xiao et al. 2017; Ornelas-Ayala et al. 2023). On the other hand, we identified changes in the expression of some of the components of the TrxG complex, which acts antagonistically with PcG. The composition of TrxG complex is variable and includes different protein types, such as histone methyltransferases (HMTs), histone demethylases, chromatin ATP-dependent remodelers, COMPASS proteins (COMplex Proteins ASsociated with SET1), among other accessory proteins. Our study showed that *mir394a mir394b* mutant plants exhibit altered expression of *ATX1* and *ATXR3*, coding for two SET-domain-containing proteins that have been shown to play roles in the regulation of flowering time and in the formation, placement, and identity of flower organs, as well as in the regulation of flowering genes (Saleh et al. 2008; Fromm and Avramova 2014; Liu et al. 2017; Xing et al. 2018). In our study we also observed upregulation of *TRO* and *WDR5a* genes, components of COMPASS-like complex. These genes are also involved in the deposition of H3K4me3 activating marks and have been shown to be involved in the regulation of floral transition and in the regulation of the floral repressor *MAF4,* which was also found to be differentially expressed in *mir394a mir394b* mutant plants, as well as *MAF5* and *NF-Y* transcription factors *YB10, C1* and *C4*, all shown to participate in the regulation of flowering time in *Arabidopsis* (Kumimoto et al. 2008; Jiang et al. 2011; Gu et al. 2013; Siriwardana et al. 2016; Zhao et al. 2017; Zhao et al. 2018; Ortuño-Miquel et al. 2019).

Taken together the results regarding chromatin remodeling factors, our study indicates that miR394 could play a role in the regulation of the deposition of activating marks on histones. Considering that the chromatin modifications typically act on a large number of genes, the observed changes in the expression of these factors are likely to impact on the expression of several of the differential expressed genes detected in this study, including the floral integrator genes, the floral meristem identity genes and those involved in floral organ development mentioned above, since this kind of regulation is widely spread in the gene regulatory networks involved in the different aspects of the flowering process in *Arabidopsis* and other species (Pien et al. 2008; Cartagena et al. 2008; Jiang et al. 2009; Berr et al. 2015; Liu et al. 2017; Cheng et al. 2017b; Jiang et al. 2018; Zhao et al. 2019; Pelayo et al. 2021; Qi et al. 2022).

Moreover, the alterations in these chromatin remodeling factors could be also affecting several of the biological processes identified in the GO term enrichment analysis presented here. Even though our downstream analysis focused on *Arabidopsis* flowering, we showed several enriched processes that suggest a role for miR394 in other aspects of plant growth and development. This is consistent with the different publications that have contributed to study the many roles of miR394 in leaf morphology, shoot apical meristem development, axillary bud initiation, response to biotic and abiotic stress, among others (Song et al. 2012; Knauer et al. 2013; Song et al. 2013; Jin and Wu 2015; Song et al. 2015; Song et al. 2016; Chand et al. 2016; Li et al. 2018; Tian et al. 2018; Tian et al. 2018; Qu et al. 2019; Kumar et al. 2019; Geng et al. 2021; Zhang et al. 2021; Bernardi et al. 2022; Lu et al. 2023; Li et al. 2024; Sun et al. 2024; Liu et al. 2024).

Finally, our transcriptomic analysis is complemented here with a detailed histological analysis of *MIR394A* and *MIR394B* reporter lines, which allowed us to establish that both genes are broadly expressed during *Arabidopsis* vegetative development, and we focused especially in the different parts of young and mature flowers. The patterns of staining indicate that both genes have overlapping areas of activity, but also specific domains such as stigmas and anthers for *pMIR394B*, but not for *pMIR394A*, or leaf margins for *pMIR394A* and leaf veins for *pMIR394B*. This level of specificity for the expression of these genes could be related to the roles of miR394 in the regulation of gene expression during plant development and in the fine tuning of the regulation of flowering time and flower development.

We had previously shown that miR394 regulation of flowering time did not involve LCR, the only characterized target of this miRNA. Also, using degradome analysis of several plant tissue samples from *Arabidopsis* (roots, leaves from different stages of development, flowers and seedlings) we also established that no additional targets regulated by transcript cleavage could be detected in *Arabidopsis* (Bernardi et al. 2022). Thus, the underlaying molecular mechanism by which miR394 regulates the chromatin remodeling factors, and the flowering genes detected in this study, remains to be determined, since this transcriptomic analysis is an insufficient approach to identify targets not regulated by transcript cleavage. The molecular mechanism might involve regulation of target transcripts by translation inhibition or sequestration of miR394 by an endogenous target mimic (eTM), as was reported for miR172 and the regulation of AP2 in the regulation of flowering time in *Arabidopsis* (Aukerman and Sakai 2003) or miR394 and the lncRNA40787 acting as an eTM in tomato to play a role in tomato resistance to late blight (Zhang et al. 2021). Therefore, the identification of these kind of targets is beyond the possibilities derived from this transcriptomic approach. Nonetheless, the results presented here reveal novel findings of the regulatory networks affected in *mir394a mir394b* mutant plants to give insight into the biological processes regulated by miR394 in *Arabidopsis thaliana*

## Acknowledgements

Authors would like to thank Marcos Reyes and Mauro Alisio, CPA members of the Argentinean Consejo Nacional de Investigaciones Científicas y Técnicas (CONICET) for collaboration with plant care and maintenance. MD, AV and AR are members of the Researcher Career of the Consejo Nacional de Investigaciones Científicas y Técnicas (CONICET) and FB is a CONICET doctoral fellow.

## Author contributions

MD was responsible for study conception and design. Material preparation, data collection and analysis were performed by FB, YB, AR and AV. The first draft of the manuscript was written by MD and FB, and all authors commented on previous versions of the manuscript. All authors read and approved the final manuscript.

## Data availability

All data generated or analyzed during this study are included in this published article and its supplementary information files.

## Declarations

Competing interests: The authors have no relevant financial or nonfinancial interests to disclose.

## Supplementary files

**Fig. S1:** Expression of *MIRAST* (AT1G09607) according to Klepikova Arabidopsis atlas.

**Fig. S2:** Mapping of small RNAs libraries from flower buds, mature leaves, seedlings and young roots.

Additional File 1: Primers used

Additional File 2: Sequence information for constructs

Additional File 3: RNAseq mapping statistics

Additional File 4: Analysis of DEG

Additional File 5: GO term enrichment

Additional File 6: Expression of genes from flowering pathways

Additional File 7: Expression of genes involved in flower development and chromatin remodeling

