## Supplementary figures and images for "Transcriptomic insights into the role of miR394 in the regulation of flowering time in *Arabidopsis thaliana*"

### Figure S1

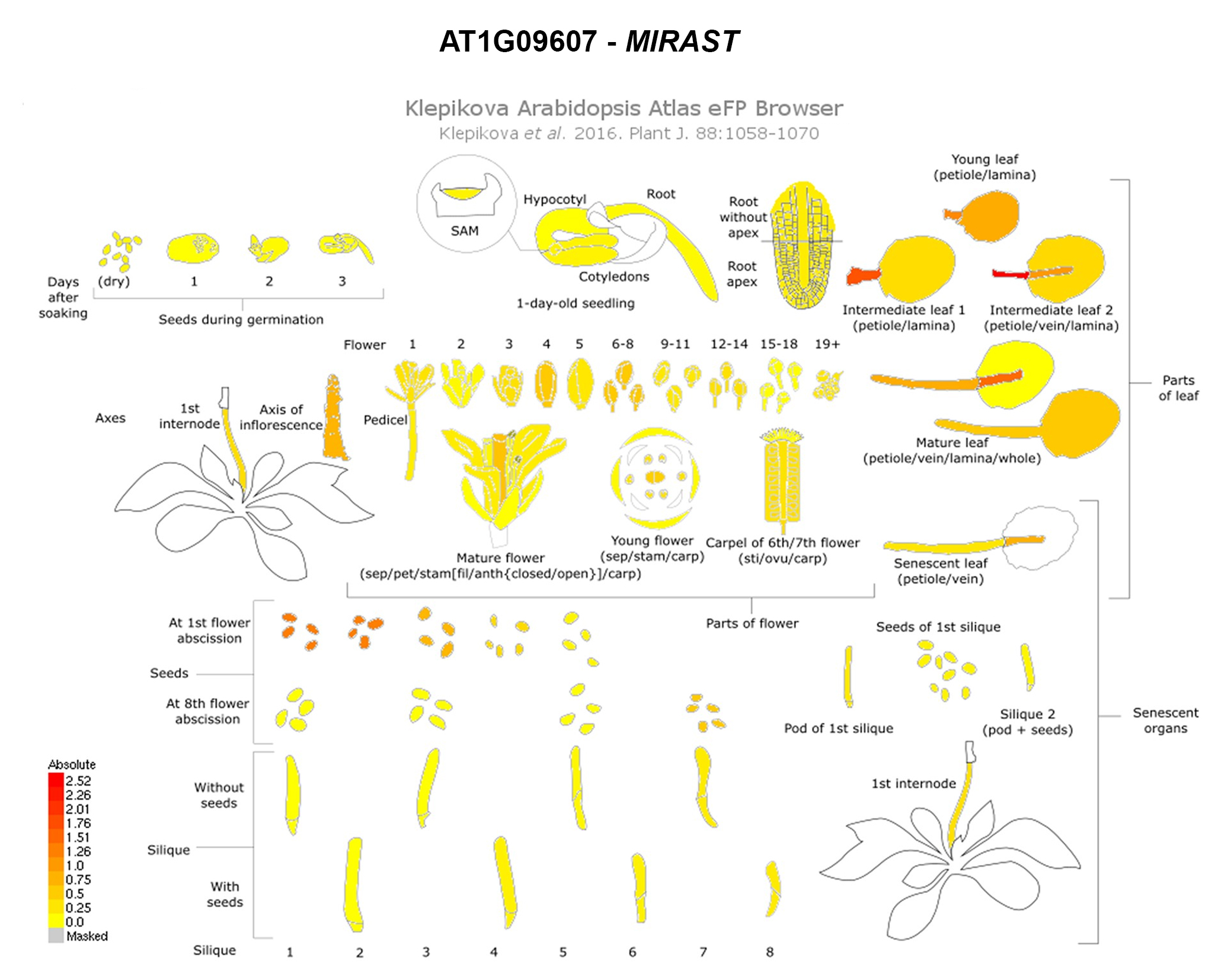

### Figure S2

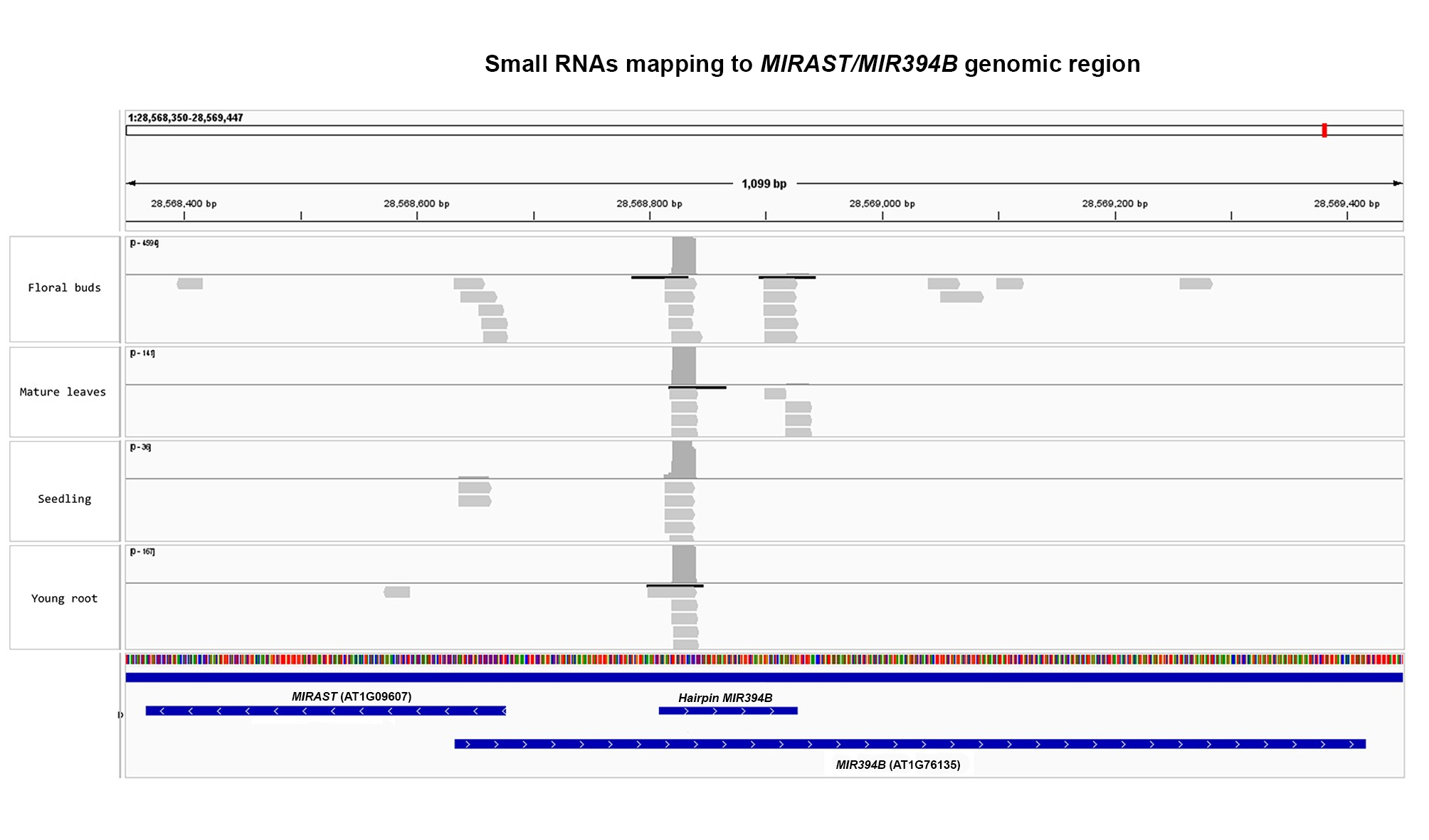
